# Aggregation-prone peptides modulate interferon gamma functionality in naturally occurring protein nanoparticles

**DOI:** 10.1101/510636

**Authors:** José Vicente Carratalá, Olivia Cano-Garrido, Julieta Sánchez, Cristina Membrado, Eudald Pérez, Oscar Conchillo-Solé, Xavier Daura, Alejandro Sánchez-Chardi, Antonio Villaverde, Anna Arís, Elena Garcia-Fruitós, Neus Ferrer-Miralles

## Abstract

Efficient protocols for the production of recombinant proteins are indispensable for the development of the biopharmaceutical sector. Approximately 400 recombinant protein-based biopharmaceuticals have been approved in recent decades, with steady growth projected in the coming years. During the expression of a heterologous gene, the protein quality control network is overcome by the disruption in protein homeostasis, leading to protein aggregation. This phenomenon has been described in all expression systems analyzed to date, including prokaryotic and eukaryotic host cells. These protein aggregates have long been considered inert protein clumps devoid of biological activity and their study has largely been neglected. However, in recent years, the classic view of protein aggregates has completely changed with the recognition that these aggregates are a valuable source of functional recombinant proteins. In this study, bovine interferon-gamma (rBoIFN-γ) was engineered to enhance the formation of protein aggregates by the addition of aggregation-prone peptides (APPs) in the generally recognized as safe (GRAS) bacterial *Lactococcus lactis* expression system. The L6K2, HALRU and CYOB peptides were selected to assess their intrinsic aggregation capability to nucleate protein aggregation. These APPs enhanced the tendency of the resulting protein to aggregate at the expense of the total protein yield. However, fine physicochemical characterization of the resulting intracellular protein nanoparticles (NPs), the protein released from these protein NPs, and the protein purified from the soluble cell fraction indicated that the compactability of protein conformations is directly related to the biological activity of variants of IFN-γ, which is used here as a model protein with therapeutic potential.

**Importance:** The demand for recombinant proteins in the pharmaceutical industry is steadily increasing. Emerging novel protein formulations, including naturally occurring protein NPs, might be an alternative to soluble variants for fine analysis at the biophysical level. Such analyses are important to address safety about biological molecules.

This study analyzes the effect of aggregation-prone peptides (APPs) on the improvement of the production of naturally occurring protein nanoparticles (NPs) of interferon gamma (IFN-γ) in the generally recognized as safe (GRAS) *Lactococcus lactis* expression system. In addition, the fine physico-chemical characterization of the resulting proteins, either obtained from the soluble or insoluble cell fractions, indicates that the selected engineered proteins embedded in the protein NPs show higher compactability than their soluble protein counterparts. Conformational compactability is directly related to the biological performance of the recombinant IFN-γ.

## Introduction

The efficient production and purification of recombinant proteins in a wide range of expression hosts has driven the launch of a large number of biopharmaceutical products (1,2). One of the most-studied expression systems is *Escherichia coli (E. coli)*, which is also one of the most-used gene expression systems for biopharmaceutical products (2,3). The *E. coli* expression system is simple, fast, robust, productive and scalable and has a wide variety of available expression vectors. In addition, its physiology has been exhaustively studied, thus allowing the optimization of production processes (4). However, prokaryotes show certain limitations, mainly the absence or limited ability to incorporate posttranslational modifications in recombinant proteins. In some instances, those modifications can be incorporated through the use of specific strain mutants or transformed strains with genes of interest. However, the introduction of eukaryotic glycosylations modifications in heterologous recombinant proteins is not feasible. Finally, pro-inflammatory contaminant lipopolysaccharide (LPS) components of the outer leaflet of the outer membrane of *E. coli* need to be removed from the purified protein to ensure the safety of the final product, increasing the final cost (5,6). Prokaryotic endotoxin-free expression systems are being explored to overcome this limitation, including *E. coli* LPS mutant strains and generally recognized as safe (GRAS) microorganisms, such as *Lactococcus lactis (L. lactis)*. These strains are envisiones as sound alternatives that avoid the safety concern of LPS contamination retaining the advantages of culturing prokaryotic hosts (1,7).

During recombinant gene expression, the great stress posed to the protein quality control machinery leads, in most cases, to the accumulation of the recombinant protein in aggregates that form intracellular nanoparticles. These aggregates are named inclusion bodies (IBs) in *E. coli*, IB-like nanoparticles in other prokaryotic hosts and aggresomes in eukaryotic hosts (7). The ratio of the amount of recombinant protein obtained in the soluble cell fraction (soluble version of the protein) to the amount of protein accumulated in the IBs (insoluble version of the protein), that is, protein solubility, depends on several factors. The solubility of a heterologous recombinant protein depends on the presence of hydrophobic residues on the surface of the protein (8) and its accessibility to the protein quality control machinery, among other factors (9). The most common strategies for improving the solubility of aggregation-prone proteins, include reducing the growth rate of the cell by reducing the growth temperature, adapting the medium composition or using weak promoters (4,10). In most cases, the recombinant protein is purified from the growth medium when secreted or from the soluble cell fraction when intracellularly accumulated. Therefore, the solubility of the recombinant protein is the key parameter evaluated when stablishing the production and purification processes. However, as more and more examples are reported, it has been accepted that during recombinant gene expression, the cell is subjected to high stress. In addition, in most instances, a certain fraction of the recombinant protein is trapped in insoluble aggregates that form discrete units in the expressing cells. These aggregates have been widely described in both prokaryotic and eukaryotic hosts (11–13).

Intracellular protein aggregates are dynamic and complex nanostructures with a variable content of recombinant protein (14–16). The trapped protein was formerly thought to be biologically inert due to, aberrant protein conformations or inactive partially folded species incompatible with the presence of biological activity. Thus, the recombinant protein could be often recovered, with low efficiency, from the insoluble cell fraction by in vitro denaturing/refolding processes (17). This scenario has been replicated in biotechnological research, and the main goal recombinant protein production is to maximize protein solubility during the production process.

However, in recent decades, the view of naturally occurring protein aggregates as inert material has changed completely since the detection of biologically active protein embedded in these aggregates (18–20). The classic view of protein aggregates as mere inactive folding intermediates has been transformed into the idea of heterogeneous porous multimeric structures stabilized by a scaffold of cross beta-sheet structures that contain conformers of the recombinant protein in which a spectrum of recombinant protein species containing native-like conformations are incorporated. In fact, further ground-breaking studies have suggested applications of IBs in nanomedicine. In these studies, IBs are envisioned as recombinant protein depots capable of slow release of active recombinant protein to replace specific biological activities in defective cell lines, to recover cell viability under stress conditions in cell culture, or even to target specific cell types in tumors when subcutaneously implanted in animal models (21,22). In addition, IBs have been proposed as a novel biomaterial for use in tissue engineering due to the stimulation of mechanical and physical signals induced in surrounding cells, even in the absence of cell growth-promoting protein factors in their formulation (23,24).

In this scenario, interest in the possibility of controlling the aggregation of recombinant proteins in these types of nanostructures is increasing, and several aggregation-prone peptides (APPs) have been identified for fusion with recombinant proteins to enhance the aggregation process in the producing cell (25). In this study, we selected rBoIFN-γ, as a model protein to study the effect of the addition of APPs in naturally occurring protein aggregates due to interest in this activity in biomedicine and its potential use in animal health.

Interferon-gamma (IFN-γ) is one of the biopharmaceuticals approved by the FDA under the trade name ACTIMMUNE^®^ (Horizon Pharma, Inc., USA). An Iranian biosimilar (γ-IMMUNEX, Exir Pharmaceutical Company, Iran) is distributed in Asia. IFN-γ is the sole type IFN. This cytokine is produced and secreted by different cell types and is involved in immunostimulatory processes through the activation of specific cellular pathways at the transcriptional level (26). IFN-γ secretion, by natural killer (NK) cells and antigen-presenting cells (APCs), enhances the innate immune response against detected pathogens, while T-lymphocytes are involved in the secretion of IFN-γ in the adaptive immune response (27,28). IFN-γ secretion is mainly regulated by other cytokines produced by APCs, including IL-12 and IL-18. The activity of IFN-γ depends on its interaction, as a dimer, with the IFN-γ receptor (IFNGR). IFNGR is a tetrameric complex formed by two ligand-binding proteins (IFNGR1) and two signal-transducing proteins (IFNGR2). The limiting factor in the transduction signal is the amount of IFNGR2 in the cell, which provides a secondary phenotype of response in those cells in which the amount of IFNGR2 is limiting, leading to partial activation of the IFN-γ-derived cascades. In that case, the proapoptotic signal of IFN-γ is not manifested, and affected cells are able to proliferate even in the presence of IFN-γ (26).

IFN-γ is used to prevent infectious diseases in patients suffering from chronic granulomatous disease and is also indicated to slow osteopetrosis (29). In recent years, IFN-γ has been investigated in approximately 80 clinical trials for a number of indications, mainly related to immune system disorders, infectious diseases and cancer (https://www.clinicaltrials.gov). Recombinant IFN-γ is administered either as a unique drug in these clinical trials (30–36) or in combination with other products (37–39). Therefore, the central key role of IFN-γ in immunostimulatory and immunomodulatory effects will lead to an increase in the use of this cytokine in the coming years in human health. The immunostimulatory properties of this cytokine have also been evaluated as alternative therapeutics in animal health (40). In fact, the administration of IFN-γ by intramammary infusion in productive dairy cows for mastitis treatment has been proposed as a putative strategy to reduce the spread of antibiotic resistance among zoonotic bacteria (41). Furthermore, the wide use of antibiotics in the prevention of animal diseases and growth promotion appears to be a source of resistant bacteria, and WHO and the United Nations have deployed global action against this threat to health security. Therefore, the rational use of antibiotics in addition to the development of immunostimulatory alternatives in the treatment and prevention of animal diseases may play a role in the control of antibiotic resistance.

Interestingly, recombinant products for animal health must not only be stable and safely and effectively delivered on a large scale and under standard conditions but also present obvious advantages over existing products to prove its commercial viability. In this context, approved recombinant human IFN-γ can be obtained from the *E. coli* expression system, but novel protein formulations need to be developed. In fact, in most reported studies of the expression and purification of IFN-γ, the recombinant protein is recovered from the purified IBs through extensive denaturation-refolding processes (42–44).

Given the importance of expanding the catalogue of prokaryotic expression strains to improve biophamaceutical safety, to explore the possibility of formulating recombinant proteins in alternative cost-effective formats and developing novel treatments to reduce the reliance on antibiotics, in this work, we produced recombinant bovine IFN-γ (rBoIFN-γ) in GRAS lactic acid bacteria *(L. lactis)* in the form of protein nanoparticles (NPs). We analyzed the ability of APPs fused to rBoIFN-γ to enhance the aggregation propensity of the recombinant cytokine. Furthermore, we assessed the link between the biological activities contained in protein NPs of IFN-γ variants and their physico-chemical characteristics. We determined that the activity of the IBs is related to the specific biological activity of the recombinant protein they contain, whereas the proportion of released protein is not the main factor. The data presented here illustrate the great potential of endotoxin-free protein NPs as active biomaterials to formulate, at the nanoscale level, proteins of biomedical interest.

## Results

### Production of rBoIFN-g in *L. lactis*

The IFN-γ gene of mammalian species encodes a preproprotein of 155-166 amino acids, including a signal peptide of 22-23 amino acids and a propeptide in the last few residues, rendering a mature protein of 15.6 to 17 kDa. Heterogeneity of the C-terminus has been described, giving rise to variants of human IFN-γ ending at residues 150, 160 or 161 (45). Human IFN-γ is usually produced in the *E. coli* expression system and is purified from IBs by using denaturing/refolding methodologies since the soluble version of the protein is proteolyzed (42–44). The same strategy has been used for mouse IFN-γ (46). In other approaches, recombinant proteins of bovine and ovine origin are obtained from the soluble cell fraction of *E. coli* and *Corynebacterium glutamicum* (47,48). We selected the rBoIFN-γ as a protein model to explore in detail different protein formats obtained from cell factories with the aim of opening up new opportunities for novel protein platforms.

In the present work, the mature bovine IFN-γ protein (residues 24 to 166) was produced in *L. lactis* with different APPs to evaluate the ability of the peptides to increase protein aggregation and to analyze the biological activity retained in the naturally occurring protein aggregates. To improve gene expression, the DNA sequence of the recombinant gene was codon optimized for the *L. lactis* expression system. Three peptides, CYOB, HALRU and L6K2, were selected based on their predicted aggregation propensity (Table 1 and Fig. 1a). AGGRESCAN was used to identify aggregation-prone segments in proteins deposited in the Disprot protein database version v6.02 (49). CYOB was selected as the peptide displaying the highest hot spot area (HSA). HALRU showed a high normalized hot spot area (NHSA) and average aggregation-propensity hot spot (a^4^vAHS) while maintaining a significantly high HSA value relative to the other identified peptides. Finally, L6K2 was previously identified as a surfactant-like peptide with the ability to enhance the aggregation propensity of several proteins (50). In the analysis, this peptide exhibited high NHSA and a^4^vAHS values despite having shorter sequence. A linker with a predicted random coil conformation was positioned between the IFN-γ and APP as previously described (50).

**FIG 1.**
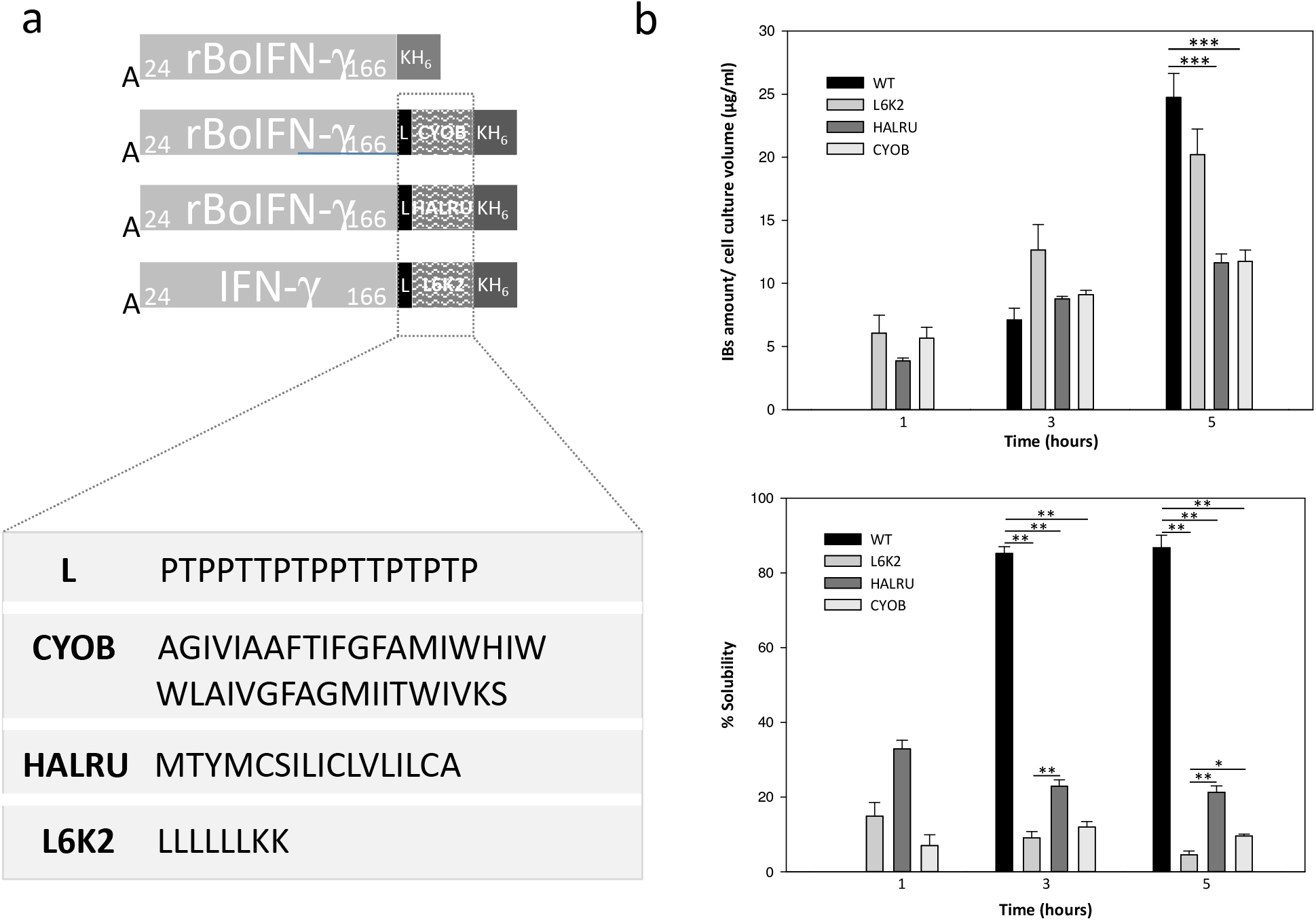
(a) IFN-γ constructs produced in *E. coli* and *L. lactis*. Residues of the IFN-γ protein are depicted in the corresponding light grey rectangles. APPs are indicated as wavy pattern boxes. The amino acid sequences of the APPs and the linker (black rectangles) between bovine IFN-γ and the APPs are shown below the drawings. The H_6_-tag fused to the C-terminus of all constructs is shown in dark grey. (b) Quantification of the production of IFN-γ in IB-like nanoparticles in *L. lactis* and solubility of IFN-γ in *L. lactis*. Significant results are shown as * *p* ≤ 0.05 and ** *p* ≤ 0.005.

**Table 1.** Selection of APPs from predictions of “hot spots (HS)” of aggregation in polypeptides by AGGRESCAN (76). CYOB: Cytochrome bo_3_ ubiquinol oxidase subunit 1 from *E. coli*, HALRU: Aragonite protein AP7. NA: Not applicable. HS: hot spot. HSA: hot spot area. NHSA: normalized HSA. a^4^vAHS: average aggregation-propensity in each HS.

| Name | UNIPROT | HS<br>region | HS<br>size | Sequence | HSA | NHSA | a<sup>4</sup>vAHS | Ref |
| --- | --- | --- | --- | --- | --- | --- | --- | --- |
| CYOB | P0ABI8 | 591-629 | 39 | AGIVIAAFSTIFGFAMIWHI<br>WWLAIVGFAGMIITWIVKS | 30.696 | 0.787 | 0.767 | This study |
| HALRU | Q9BP37 | 1-17 | 17 | MTYMC SILICLV LILCA | 15.842 | 0.932 | 0.904 | This study |
| L6K2 | NA | 1-6 | 6 | LLLLLLKK | 6.211 | 1.035 | 0.949 | (50) |

In *L. lactis*, most of the protein is detected in the soluble cell fraction in the absence of any of the APP (Fig. 1b). This observation is in agreement with previous results for the expression of the natural DNA sequence of the bovine IFN-γ gene in *E. coli* in which His-tagged rBoIFN-γ was purified by affinity chromatography from the soluble cell fraction (48). The presence of the APPs in the recombinant protein caused a noticeable shift of the final products toward the insoluble cell fraction, as expected (Fig. 1b). The APP resulting in the highest aggregation tendency was the L6K2 peptide. In addition, the presence of an APP tag also had a negative effect on the total recombinant protein produced in the cell. This negative effect was maximal at 3 h, when protein levels of 13.82 ± 2.01 μg/ml, 11.38 ± 0.36 μg/ml, and 10.36 ± 0.45 μg/ml were observed for the IFN-γ variants fused with the L6K2, HALRU and CYOB peptides, respectively, compared with 211.99 ± 51.46 μg/ml for wild type IFN-γ. Therefore, the best APP in terms of aggregation propensity and protein yield in the insoluble cell fraction, corresponded to the IFN-γ L6K2 formulation. Surprisingly, the performance of this surfactant-like peptide exceeded the predicted aggregation-prone capabilities of CYOB and HALRU peptides (Table 1).

### Nanoarchitectonic characterization of protein nanoparticles

The morphometry of purified protein NPs of the rBoIFN-γ variants was examined by field emission scanning electron microscopy (FESEM; Fig. 2a). The images revealed the presence of multimeric complexes comprising discrete NPs in addition to isolated protein NPs (inset Fig. 2a). First, the NPs were similar to bovine IFN-γ protein NPs obtained previously in this expression system (7). Z potential (ZP) measurements showed that all of the NPs presented negatively charged surfaces with negative values ranging from −38 to −28 mV (Fig. 2b), indicating the stability of the NP suspension. The higher values of ZP obtained for the IFN-γ variants provide information about particle stability, as NPs displaying higher ZP values (higher than +30 mV or lower than −30 mV) exhibit increased stability due to greater electrostatic repulsion between particles (51).

**FIG 2.**
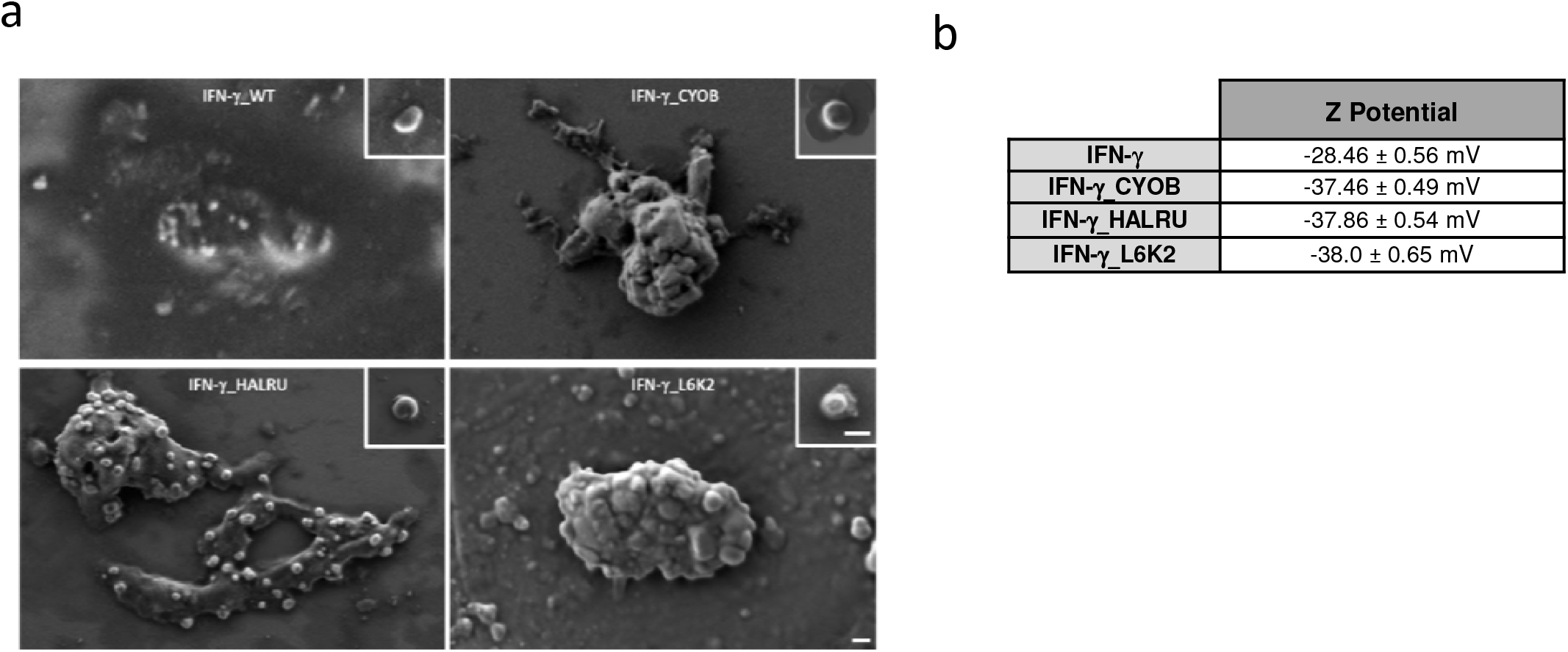
(a) Ultrastructural characterization by FESEM of protein aggregates and purified protein nanoparticles of rBoIFN_γ, rBoIFN-γ_CYOB, rBoIFN-γ_HARLU and rBoIFN-γ_L6K2. Scale bars correspond to 200 nm. (b) Z potential of purified protein nanoparticles.

### Biological activity of soluble IFN-γ and nanoparticles of IFN-γ

The activity of IFN-γ is usually determined by an antiviral assay (52). This assay must be performed in facilities with an appropriate biosafety level, and viral stocks have to be maintained over time. Therefore, alternative assays have been developed to simplify the procedure. One approach to evaluate IFN-γ activity mediated by IFN-γ-receptor binding is the detection of kynurenine. The antiproliferative activity of IFN-γ in this assay is related to the induction of the expression of the indoleamine-2, 3-dioxygenase (IDO) gene, which is the first and rate-limiting enzyme in tryptophan catabolism. IDO catalyzes oxidative cleavage of tryptophan to N-formylkynurenine. Following a hydrolysis step, the latter is transformed into kynurenine by Ehrlich’s reagent, giving a yellow-colored compound absorbing at 490 nm (53). To validate the assay, the activity of three soluble rBoIFN-γ proteins was tested by using the kynurenine detection method (Fig. 3a). rBoIFN-γ_Std corresponded to a commercially available mixture of bovine IFN-γ Gln24-Thr166 and Gln24-Arg162, both with an N-terminal Met (R&D Systems). This protein preparation exhibited the lowest dissociation constant (K_D_) among the proteins purified from the soluble cell fraction (Fig. 3a). The difference in this parameter with in-house IFN-γ produced in *E. coli* (rBoIFN-γ_E. *coli*) was related to the Gln24-Arg162 variant, which was not included in the protein preparation. However, this difference may also be attributable to other variables, such as the buffer composition of the protein stock or the folding abilities of the transcribed mRNAs from the two genes encoding the same protein but with different codon usages in their sequences (54). The protein obtained from the *L. lactis* expression system displayed less activity, which may be to differences in the way *L. lactis* and *E. coli* cope with gene codon usage that favor the specific activity of the protein obtained from *E. coli*. Other variables may apply, such as the induction temperature (30 °C for *L. lactis* and 37 °C for *E. coli)* or even the presence of a His-tag purification motif in the recombinant variants which is not present in the commercial version of the protein. Alternatively, the growth medium used in the protein production experiments might result in different growth rates and folding efficiencies (55,56). To determine the biological activity contained in the IFN-γ protein NPs produced in *L. lactis*, similar experiments were performed by incubating the bovine cells with increasing amounts of rBoIFN-γ NPs, that is, protein NPs purified from the insoluble cell fraction (Fig. 3b). The results showed that all cells were able to elicit responses to the presence of the protein NPs, and the IFN-γ_L6K2 formulation displayed the highest initial rate and kynurenine production. The addition of HALRU and CYOB APP to IFN-γ had a moderate effect on the cell response of the. As shown in Fig. 3a and Fig. 3b, the experiments were performed at the same time with the same stock of cells and conditions. We wondered why the sample corresponding to protein NPs of IFN-γ_L6K2 had the highest activity and initial rate, even compared with commercial IFN-γ. Consistent with this observation, a previous analysis of the activity of recombinant β-galactosidase in the form of protein NPs revealed higher specific activity than the corresponding soluble version of the protein (18). However, these protein samples have not been characterized in detail. The activity displayed by IBs produced in *E. coli* has been attributed to the release of a spectrum of conformers of the recombinant protein, which leaves a scaffold that is resistant to proteolysis and has an extensive cross-beta-pleated sheet conformation (57,58). In fact, in the case of protein NPs of rBoIFN-γ produced in *L. lactis*, 30-40 % of the material is resistant to proteolysis, indicating that the protein NPs obtained in this expression system follow the same principles as the *E. coli* system (7). Therefore, the activities displayed by the protein NPs are due to the partial release of the IFN-γ that forms part of the macromolecular complex (22). The ability of the protein NPs to release protein was evaluated by incubating them in PBS for 96 h to emulate the protein release conditions established during the biological activity assay of the protein NPs (see the experimental design used to obtain the different protein samples in Figure 3c). Release of 52.67 %, 5.30 %, 0.42 % and 0.46 % was observed for IFN-γ, IFN-γ_L6K2, IFN-γ_HALRU and IFN-γ_CYOB, respectively. The highest activity among all samples shown in Fig. 3a and Fig. 3b corresponds to IFN-γ_L6K2 NPs, which have a limited ability to release recombinant protein. Thus, an activity assay was performed to analyze the specific activity of the proteins released from the protein NPs compared with soluble proteins obtained directly from the soluble cell fraction (Fig. 3d). The results showed that the maximal specific activity corresponded to the IFN-γ_L6K2 protein released from NPs. In addition, the comparison of the specific activity of the rBoIFN-γ protein produced in *L. lactis* and purified from the soluble cell fraction with that of the corresponding protein released from NPs showed that the released protein elicitsed better conformational performance (compare the second and last bars in Fig. 3d).

**FIG 3.**
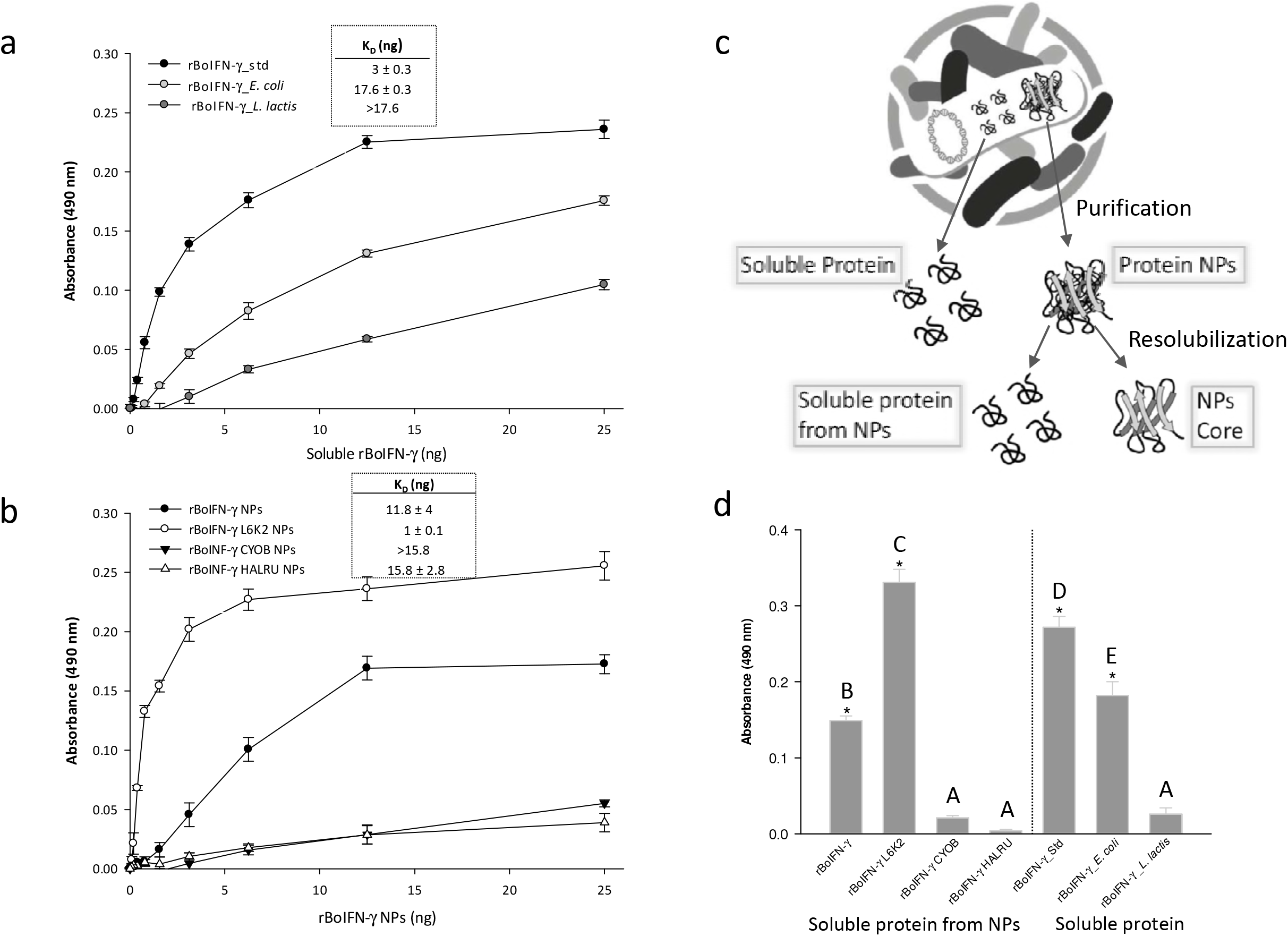
Kynurenine levels measured by absorbance at 490 nm after treatment of EBTr cells for 96 h with increasing amounts of rBoIFN-γ from different origins. (a) Soluble rBoIFN-γ produced in the indicated expression system. (b) Protein nanoparticles of rBoIFN-γ produced in *L. lactis*. The Kd values are indicated in the plot. (c) Schematic representation of the protein samples used in the activity assays: soluble protein obtained from the soluble cell fraction, protein NPs purified from the insoluble cell fraction, soluble protein obtained from the protein NPs, and the NP core after a resolubilization procedure. (d) A 3 ng quantity of rBoIFN-γ protein was obtained from solubilization of protein NPs or purified rBoIFN-γ from the soluble cell fraction as indicated. Different letters depict differences between proteins (*p* < 0.001) except rBoIFN-γ from protein NPs and rBoIFN-γ_E (p = 0.024).

### Physicochemical characterization of soluble IFN-γ and nanoparticles of IFN-γ

To further analyze the protein in different formats, DLS measurements were performed (Fig. 4a-4d). The commercial bovine IFN-γ exhibited a peak with a maximum at 7.6 nm, was quite similar to the peak at 6.13 nm for the in-house version of IFN-γ produced in *L. lactis*. This configuration might correspond to the dimeric form of the cytokine. However, the IFN-γ obtained in-house in *E. coli* showed a tendency towards a larger size. Therefore, the specific activity of the different rBoIFN-γ formats is not directly linked to the dimeric configuration, which is the functional conformation when binding to the cell receptor, and some other variables might be involved. When analyzing the size of the purified NPs, a peak above 1000 nm was detected, which is above the upper sensitivity limit of the equipment (Fig. 4b). The NPs were clustered in higher-order complexes from monomeric versions of 200 nm (Fig. 2a). All samples exhibited the same profile. After solubilization of the protein embedded in the NPs, the size of the remaining material remained above 1000 nm since the scaffold of the NPs retained the overall structure after the protein was released (Fig. 4c). The released protein showed a narrow dispersion ranging from the dimeric size of the protein identified in the samples obtained from the commercial IFN-γ or the soluble version purified from *L. lactis* detected in the upper panel of Fig. 4a (Fig. 4d). In addition, the polydispersity index (PI) of these samples was higher than that of the soluble IFN-γ versions. The PI corresponds to an estimate of the width of the distribution, and the higher values of PI are in accordance with the data showing the presence of a pool of conformers in the folding of recombinant proteins when the proteins are produced in the cell. By contrast, in the protein versions purified from the soluble cell fraction, the downstream process selects only a narrow collection of conformers, indicating that the protein obtained during solubilization assays from protein NPs is more representative of the diversity in conformations of a single protein that are produced in the expression system.

**FIG 4.**
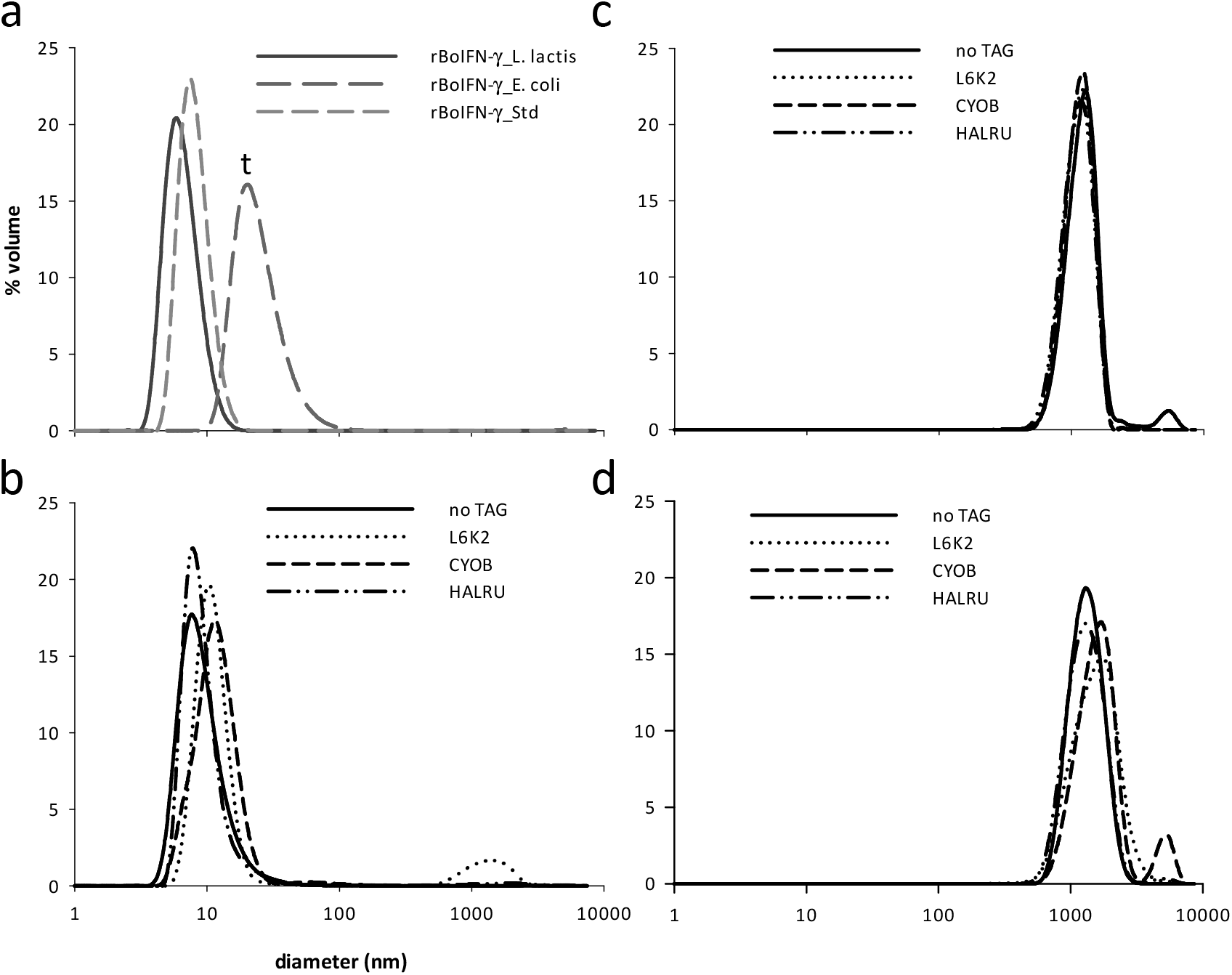
Recombinant IFN-γ sizes in different supramolecular arrangements (purified soluble IFN-γ and INF-γ IBs). (a) Soluble rBoIFN-γ from different origins: commercial rBoIFN-γ, inhouse rBoIFN-γ from *E. coli* and *L. lactis*. (b) rBoIFN-γ IBs produced in *L. lactis*. (c) Scaffold of rBoIFN-γ IBs incubated for 96 hours at 37 °C and (d) resolubilized rBoIFN-γ from IBs after interferon release (the mean size and polydispersity index are indicated in brackets). The average size data of the soluble proteins were analyzed by one-way ANOVA (^t^ corresponds to *P* < 0.07).

To further analyze the link between the physico-chemical properties and the specific activity of the proteins, the fluorescence emission of Trp was recorded. Each fluorescence emission spectrum was transformed into a center of spectral mass (CSM) value. This parameter is related to the relative exposure of the Trp to the protein environment. The maximum red-shift in the CSM of the Trp spectrum is compatible with large solvent accessibility (59–61). By contrast, the blue shift in the CSM corresponds to a Trp hidden in a more hydrophobic milieu (62). The mature form of BoIFN-γ has a unique Trp. This Trp residue is partially buried in the 3D structure of the protein (PDB 1D9C) (63) and is not involved in either monomer or in cytokine-receptor interactions, as shown in the 3D structure of the human tetrameric complex of the cytokine dimer with the receptor (PDB 1FG9) (64). A remarkable aspect of the intrinsic fluorescence analysis is that all of the rBoIFN-γ variants within the NPs or after solubilization from the protein NPs exhibited lower CSM values than the samples obtained from the soluble protein (Table 2). These results suggest that the protein forming part of the NPs and the protein solubilized from the aggregates have a more compact conformation than the soluble version. The differences between the soluble IFN-γ versions are due to differences in protein origin, sequence and size. The most efficient IFN-γ soluble version corresponded to the commercial IFN-γ, which had the lowest CMS due to its highly compacted structure. The proteins obtained from the soluble fraction of *E. coli* and *L. lactis* exhibited higher CMS values than the commercial protein. These differences might be related to the distinct sizes detected (Fig. 4a). The in-house rBoIFN-γ_E. *coli* was approximately three times larger than the same NPs produced in *L lactis*, indicating that the Trp residue sensed a more polar environment compared with the *L lactis* form (Table 2). Interestingly, for the protein originating from the particulate form, a blue shift was observed comparde with the soluble versions, and the CSM increased as itwa resolubilized (lines 2 and 4 of Table 2), at least in versions with a percentage of release above 5 %. Interestingly, the CSM value of the resolubilized rBoIFN-γ_*L. lactis* protein sample did not reach that of the soluble counterpart (lines 3 and 4 of Table 2). When the APPs were incorporated in the engineered protein constructs, the resolubilized proteins showed a decrease in the CSM values compared with the protein NPs samples (lines 5 and 7). This behavior suggests a possible self-arrangement of the tag within the protein that could replace water molecules and increase the hydrophobicity of the Trp environment. The CYOB construct (line 6 of Table 2) required a specific analysis as this tag contributes five additional Trp residues to the whole protein structure. In this case, the resolubilized protein spectrum exhibited a modest red shift (higher CMS value) compared with the particulate form, indicating that the resolubilization process exposed some of the Trp residues to a hydrophilic environment. The CSM values of the CYOB and HALRU protein NPs remained unaltered after resolubilization (Table 2, lines 6 and 7). These data are in accordance with the higher stability of the particulate forms, which exhibited low levels of protein release.

**Table 2.** Center of spectral mass (CSM) of IFN-γ protein preparations in soluble formats or in protein NPs analyzed before and after the resolubilization protocol.

| | | Soluble rBoIFN-γ | rBoIFN-γ NPs | | Soluble rBoIFN-γ<br>from NPs |
| --- | --- | --- | --- | --- | --- |
|  |  | CSM (nm) |  |  |  |
|  |  | Soluble | Protein NPs | NP core | Resolubilized |
| 1 | rBoINF-γ_Std | 356.5 | | | |
| 2 | rBoIFN-γ_ *E. coli* | 358.1 | | | |
| 3 | rBoIFN-γ_ *L. lactis* | 357.4 | | | |
| 4 | rBoIFN-γ_ *L. lactis* | | 352.4 | 353.6 | 354.2 |
| 5 | rBoIFN-γ_L6K2_ *L. lactis* | | 352.7 | 353.6 | 351.7 |
| 6 | rBoIFN-γ_CYOB_ *L. lactis* | | 353.1 | 353.1 | 354 |
| 7 | rBoIFNγ_ HALRU_ *L. lactis* | | 354.2 | 354.2 | 352 |

The NP form of IFN-γ also favored the specific activity (insets, Fig. 3, *L. lactis* and nontagged rBoIFN). This phenomenon is not only due to more active conformation of the protein (Fig. 4 and Table 2, line 3 vs line 4) (65) but also to the heterogeneous distribution of the protein and the ability of the protein NPs to increase the effective concentration of protein in the proximity of the receptor. Moreover, the formulation containing L6K2 was the most efficient, even compared with the commercial protein. Resolubilization clearly conferred the most active and altered conformation of the protein without the tag. Although a low percentage of protein was released from the NPs containing L6K2, at least in the physiological buffer PBS, this released protein seems to be sufficient to surpass the activity of the released protein without a tag (Fig.3).

Another interesting aspect is the effect of the size of the tag on the structure-function of the protein. The incorporation of a tag larger than 17 aa beyond the linker (Fig. 1) could generate steric problems preventing the interaction of tagged IFN-γ with the receptor. The larger the tag, the more negative the effect on the activity. As shown in Fig. 1, L6K2 is only 8 aa, compared with 17 aa for HALRU and 38 aa for CYOB. The short size of the L6K2 tag migth reduce the difficulty of the interaction between L6K2-IFN-γ and the receptor compared with the longer IFN-γ tags.

## Discussion

The production and purification of recombinant proteins is a technology of expanding interest in research and in biopharmaceutical, and industrial applications. More than 400 protein-based pharmaceuticals have been approved, with a steady increase over the last decade of this type of NBE (1,2). The precise analysis of recombinant proteins is important due to safety concerns, and specific regulatory guidelines were redesigned for protein-based pharmaceuticals after the TGN1412 clinical trial (66,67). Therefore, the physicochemical characterization of recombinant proteins needs to be further developed, and technical approaches to obtain reliable data on the quality of the final product are needed. In this work, the biological activities of three versions of the same protein produced in two different LPS-free expression systems, *ClearColi* and *L. lactis* indicated that the production process imposed conformational restrictions on the recombinant products. For genes with the same nucleotide sequence (*L. lactis* codon-optimized rBoIFN-γ gene) but expressed in different expression systems (Fig. 3a; compare the curves of rBoIFN-γ_E. *coli* and rBoIFN-γ_*L. lactis*), the biological activity of the resulting proteins differed by an order of magnitude. However, even in the same expression system, the nucleotide sequence of the gene was also a factor influencing the quality of the purified protein (Fig. 3a, compare the curves of rBoIFN-γ_E. *coli* and rBoIFN-γ_Std, which correspond to the *L. lactis* codon-optimized or natural *Bos taurus* gene nucleotide sequences, respectively). Those results might be related to the ability of the different expression systems to cope with the overexpression of the recombinant protein. The codon usage in an expression system can affect the translation profile and hence the biological activity of the final product (54,68). In addition, the secondary structure of the resulting mRNA seems to affect the efficiency of translation and mRNA degradation (69,70). The production of rBoIFN-γ protein in *E. coli* was negligible (data not shown) when using an *E. coli* codon-optimized rBoIFN-γ gene, whereas protein production was achieved when the *L. lactis* codon-optimized gene was used, as shown in the present study. The DLS analyses of the recombinant proteins indicated that the rBoIFN-γ produced in *E. coli* from the *Bos taurus* natural nucleotide sequence and in the *L. lactis* expression system from the *L. lactis* codon-optimized sequence were similar in size, but the *L. lactis* codon-optimized gene expressed in *E. coli* resulted in a protein product of larger size (Fig. 4), confirming the effects of the expression system and codon usage on the quality of the final product. Differences in the compactability of protein samples with the same amino acid sequence were also observed (Table 2).

In this context, the addition of APPs to the rBoIFN-γ protein improved the aggregation profile of the produced protein (Fig. 1b). However, the presence of this type of peptides had a negative effect on the overall production of the protein and, in the case of HALRU and CYOB, a huge impact on biological activity (Fig. 3). Therefore, AGGRESCAN software is able to predict the propensity to aggregate of the resulting APP-containing recombinant IFN-γ and is a reliable tool for analyzing solubility performance in the design of recombinant genes. The size of the APP is an important parameter, and as a general rule, the longer the aggregation peptide, the higher the HSA. However, small peptides with a discrete HSA but high normalized and average HSA have also been reported to enhance the aggregation of the accompanying proteins, as in the case of L6K2 (50). Surprisingly, the L6K2 peptide also leads to the production of highly active protein in the protein NPs. The detailed physicochemical characterization of the protein samples revealed higher compactability of the protein conformations in the protein samples with the highest biological activity. The protein peaks of the soluble protein samples indicated that the purification strategy of a protein selects for a narrower distribution of protein conformations compared with protein samples recovered from naturally occurring protein NPs (compare the PDI of the samples obtained from the soluble cell fraction with the protein samples obtained after solubilization of protein NPs in Fig. 4).

In the recombinant protein production platform, the general consensus for improving protein yield is to improve the solubility of the protein. Consequently, the major established strategies for producing recombinant protein involve improving solubility by including folding modulators or reducing the growth rate. However, solubility and conformational quality are not necessarily coincident parameters (71). The functionalities of the protein obtained from the soluble cell fraction or the protein NPs of rBoIFN-γ_*L. lactis* in the present work were in accordance with these previous findings, as the protein obtained from the soluble cell fraction was less active than that recovered from the protein NPs. The compactabilities of the conformations of these proteins were in agreement with their dissimilar biological activity. The results obtained in this study are in accordance with previous analyses indicating that the compactability of protein conformations is a key parameter related to stability and function (72,73).

Consistent with these observations, the addition of the L6K2 APP to rBoIFN-γ_*L. lactis* improved the biological performance of the resulting conformational species retained in the corresponding protein NPs. Interestingly, this APP showed higher capability as a compactability factor compared with the other APPs tested with IFN-γ. In fact, the addition of the other two APPs to rBoIFN-γ (HALRU and CYOB) surprisingly had negative effects on the functionality of the resulting proteins, despite levels of compactability similar to that of the protein sample lacking the APPs. These sequences might have negative effects by preventing correct binding of the dimer to the cell receptor. In conclusion, the detailed physicochemical characterization of the purified recombinant proteins needs to be further developed to select a more appropriate product to optimize the quality of the recombinant product.

## Materials and methods

### Bacterial strains and plasmids

*E. coli* MC4100 (StrepR) (74) was used for cloning genes for protein production in *L. lactis. E. coli* DH5α was used for cloning genes for protein production in *E. coli. L. lactis cremoris* NZ9000 (kindly provided by INRA, Jouy-en-Josas, France; patent n° EP1141337B1), and *ClearColi*^®^ BL21(DE3) (Lucigen) were used in expression experiments for each expression system. For *L. lactis* expression vectors, IFN-γ of bovine *(Bos taurus)* origin was cloned at the *NcoI/XbaI* restriction sites of the CmR pNZ8148 plasmid (MoBiTech). The digestion products were ligated into the expression plasmid pNZ8148, and ligation product was used to transform *L. lactis* NZ9000 competent cells by electroporation (75). Electroporation was performed using a Gene Pulser from Bio-Rad with settings of 2500 V, 200 Ω and 25 μF in a pre-cooled 2-cm electroporation cuvette. The electroporated cells were then supplemented with 900 μl of M17 broth with 0. 5 % glucose and incubated for 2 h at 30 °C. The electroporation mix was centrifuged for 10 min at 10,000 x *g* at 4 °C, and the pellet was resuspended in 100-200 μl of M17 media and plated. In addition, fusions of rBpIFN-γ with APPs were constructed (rBOIFN-γ_L6K2, rBoIFN-γ_HALRU and rBoIFN-γ_CYOB). All genes were C-terminally fused to a His-tag for detection and quantification by western blot analysis. A Lys residue was included at the N-terminus of the tag for putative elimination of the tag by exopeptidases. Gene sequences were codon optimized for the *L. lactis* expression host as indicated (Geneart). For the *E. coli* expression vector pETDuet (Novagen), the *L. lactis* codon-optimized IFN-γ gene of bovine *(Bos taurus)* origin was cloned at the NcoI/HindIII restriction sites of the pETDuet plasmid. The expression vector was transformed into the *ClearColi*^®^ BL21(DE3) strain. Electroporation was performed using a Gene Pulser from Bio-Rad with settings at 2400 V, 750 Ω and 25 μF in a pre-cooled 2-cm electroporation cuvette. The electroporated cells were then supplemented with 900 μl of lysogeny broth (LB) and incubated for 1 h at 37 °C. The cells were then plated on LB-agar plates containing ampicillin (100 μg/ml) and incubated at 37 °C overnight. For all clones, in the sequence design we added an *NcoI* restriction site at the 5’ end followed by the nucleotides CA to restore the reading frame. This cloning strategy adds an Ala to the N-terminus of the protein. Therefore, the recombinant proteins were produces as the mature form of the IFN-γ (from Gln24 to Thr166 NP_776511.1) with an additional Ala at the N-terminus to restore the frameshift introduced by the *NcoI* restriction site (Fig. 1a).

### Selection of aggregation-prone peptides (APPs)

APPs were selected by scanning the Disprot v6.02 database (49)with AGGRESCAN software (76). We selected two unstructured regions from two different proteins that displayed a higher HSA. This selection was based on the assumption that APPs in solvent-exposed regions were the best candidates for the purposes of this study. Additionally, L6K2 was selected based on previous experimental results (50) after analysis with AGGRESCAN showed that this peptide had a high NHSA and high a^4^vAHS.

### Production and purification of rBoIFN-γ protein from the soluble cell fraction

Cultures of *ClearColi*^®^ BL21 (DE3) cells transformed with the plasmid containing rBoIFN-γ gene (pETDuet-rBoIFN-γ) were incubated in a shake flask at 37 °C and 250 rpm in LB medium supplemented with 100 μg/ml ampicillin. When the cultures reached an OD 550 of approximately 0.5 – 0.7, protein expression was induced by adding 1 mM IPTG (isopropyl-β-D-thiogalactopyranoside). The cultures were then incubated at 20 °C and 250 rpm overnight (for protein production). During the purification process, cells were collected by centrifugation (15 min, 6,000 x *g*, 4 °C), and proteins were released by sonication. Briefly, cells were lysed by sonication with 5 rounds of 3-min cycles and pulses of 0.5 s at 15 % amplitude (Lab Sonic ultrasonicator). *L. lactis cremoris* cells transformed with plasmids containing the rBoIFN-γ gene (pNZ8148-rBoIFN-γ, pNZ8148-rBoIFN-γ_L6K2, pNZ8148-rBoIFN-γ_HALRU and pNZ8148-rBoIFN-γ_CYOB) were incubated in a shake flask at 30 °C without shaking in M17 broth + 0.5 % glucose supplemented with 5 μg/ml chloramphenicol and 2.5 μg/ml erythromycin. When the cultures reached an OD 550 of approximately 0.4-0.6, protein expression was induced by adding 12.5 ng/ml nisin. Then, the cultures were incubated at 30 °C without shaking for the indicated time. The cells were collected by centrifugation (15 min, 6,000 x *g*, 4 °C), and proteins were released by a French press (3 rounds at 15000 PSI/machine pressure).

From this point, the same protocol was followed for both proteins. The soluble and insoluble cell fractions were separated by centrifugation (40 min, 15,000 x *g*, 4 °C), and the soluble cell fraction was filtered using a pore diameter of 0.2 μm. The recombinant protein in the soluble cell fraction was purified by immobilized metal affinity chromatography (IMAC) using a HiTrap Chelating HP 1-ml column (GE Healthcare) with an ÄKTA purifier FPLC system (GE Healthcare). The eluted proteins were then dialyzed against phosphate-buffered saline (PBS) buffer. The control protein rBoIFN-γ_Std was obtained from R&D Systems (2300-BG-025, R&D Systems).

### Production and purification of rBoIFN-γ protein nanoparticles

*L. lactis* cells transformed with expression plasmids derived from pNZ8148 were grown in M17 medium enriched with 0.5 % glucose at 30 °C without shaking. *E. coli* was grown in LB rich medium at 37 °C and 250 rpm. NP production was induced by adding 12.5 ng/ml nisin (Sigma-Aldrich) to *L. lactis* or 1 mM IPTG to *E. coli* cultures. After induction, the cultures were grown for 5 h. Antibiotics were used for plasmid maintenance at the following concentrations: chloramphenicol (5 μg/ml) and erythromycin (2.5 μg/ml) for *L. lactis* and ampicillin (100 μg/ml) and streptomycin (30 μg/ml) for *ClearColi*.

Once produced, the protein NPs were purified using the purification protocol described previously (7), including, at the beginning of the process, a mechanical disruption step by French press. The protocol was performed under sterile conditions and all incubations were carried out under agitation. The purified protein NPs were diluted 1:10 in PBS and resuspended.

### Quantitative protein analysis

The amounts of recombinant proteins produced by the expressing cells or present in NPs were quantified by denaturing SDS-PAGE as described previously (77). Bands were identified using a commercial polyclonal serum against the histidine tag (#A00186-100 Genscript) and an anti-mouse secondary antibody (#170-6516, Bio-Rad). The recombinant protein yield was estimated by comparison with a standard curve of known amounts of a purified GFP-H6 protein quantified by the Bradford assay. Quantification was performed with Quantity One software (Bio-Rad).

### Ultrastructural characterization

To characterize the morphometry (size and shape) of the NPs, microdrops of protein aggregate suspensions were deposited for 2 min on silicon wafers (Ted Pella Inc.), air-dried and observed in a nearly native state under a field emission scanning electron microscope (FESEM) Zeiss Merlin (Zeiss) operating at 1 kV. Micrographs of the NPs were acquired with a high-resolution in-lens secondary electron (SE) detector. Images were taken at magnifications ranging from 20,000x to 80,000x.

### Z potential analysis

Z potential (ZP) characterization of each kind of protein NPs was carried out using DLS equipment (Malvern Nanosizer). To prevent the electrodes from burning, the samples were prepared in deionized (MilliQ) water, a low ionic strength medium. Each sample was analyzed in triplicate.

### Determination of rBoIFN-γ biological activity in bovine cells

The different rBoIFN-γ formulations described here were analyzed by a modified kynurenine bioassay (78). Bovine fibroblast-like cells (EBTr cells), (87090202 Sigma-Aldrich) were cultured in Dulbecco’s modified Eagle’s medium (Gibco) with 10 % fetal bovine serum (FBS). Depending on cell growth, cultures were split every 2 days using ratios of 1:2 to 1:3. Before an experiment, a lower ratio was selected to obtain the maximum cell density. For activity analysis, the cells were seeded in 96-well flat-bottom microtiter plates (5000 cells per well) in Dulbecco’s modified Eagle’s medium (Gibco) supplemented with 100 μg/ml L-Trp. Serial dilutions of both, the soluble and NP forms of rBoIFNy at quantities ranging from 25 to 0.1 ng were incubated with cells for 96 h at 37 °C. Aliquots of 160 μl of the supernatants were then mixed with 10 μl of 30 % trichloroacetic acid (T6399 Sigma-Aldrich) and incubated at 50 °C for 30 min. After a centrifugation step (10 min, 600 x g), aliquots of 100 μl of the supernatants were mixed with an equal volume of 4 % w/v Ehrlich’s reagent 4-(dimethylamino) benzaldehyde (156477, Sigma-Aldrich) in glacial acetic acid (Fisher Chemical A/0360/PB15). After 10 min, the absorbance at 490 nm was measured in a conventional luminometer and VICTOR3V 1420 multilabel reader (PerkinElmer).

The absorbance vs IFN-γ quantity curves were adjusted to Eq. 1. Abs490 is the absorbance at 490 nm, which represents an indirect measurement of IFN-γ binding to the receptor, Absmax is the maximal binding of IFN-γ to the receptor, and K_D_ is the equilibrium dissociation constant. A low value of K_D_ indicates high IFN-γ affinity to the receptor.

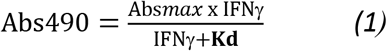

### Assay of protein solubilization from protein nanoparticles

The rBoIFN-γ protein NPs (rBoIFN-γ_L, rBoIFN-γ_L6K2, rBoIFN-γ_CYOB and rBoIFN-γ_HALRU) were solubilized in different volumes of PBS depending on their initial protein amounts. In all cases, the concentration was adjusted to 20 μg/ml. After manual agitation, every sample was incubated at 37 °C for 96 h to reproduce the conditions used during the biological activity determination. The soluble and insoluble fractions were then isolated by centrifugation (15 min, 15,000 x g). The protein amounts in the soluble fractions were quantified, and the concentrations were adjusted. The biological activity of every sample was determined at a single concentration (3 ng/μl) as described previously. The protein released from the protein NPs was resuspended in Laemmli buffer (1x) (from Laemmli buffer (4x): 1.28 g of Tris base, 8 ml of glycerol, 1.6 g of sodium dodecylsulfate (SDS), 4 ml of β-mercaptoethanol and 9.6 g of urea in a final volume of 100 ml), boiled at 98 °C for 45 min, and loaded onto SDS-polyacrylamide gel electrophoresis (10 % acrylamide) denaturing gels. Protein bands were detected by Western blot using a commercial 6xHis monoclonal antibody (631212, Clontech) and a goat anti-mouse IgG (H + L)-HRP conjugate (1706516, Bio-Rad) as the secondary antibody. Images of the membranes were obtained using the ChemiDoc Imaging System (Bio-Rad), and bands were quantified with Image Lab Software (Bio-Rad) using known concentrations of commercial rBoIFN-γ (20, 15, 10, 5, 3 and 1 ng; 2300-BG-025, R&D Systems).

### Interferon size determination

The volume size distribution of interferon γ was determined by Dynamic Light Scattering (DLS). A 60-μl aliquot (stored at −80 °C) was thawed, and the volume size distribution of each protein format was immediately determined at 633 nm (Zetasizer Nano ZS, Malvern Instruments Ltd.).

### Analysis of protein conformation by intrinsic tryptophan fluorescence

Fluorescence spectra were recorded on a Cary Eclipse spectrofluorometer (Agilent Technologies). A quartz cell with a 10-mm path length and a thermostated holder was used. The excitation and emission slits were set at 5 nm. The excitation wavelength (λ_ex_) was set at 295 nm. Emission spectra were acquired within a range from 310 to 550 nm. The protein concentration was 0.3 mg/ml in PBS. *To* evaluate conformational differences between the proteins, we applied the CSM. CSM is the weighted average of the fluorescence spectrum peak.

The CSM was calculated for each of the fluorescence emission spectra (79) according to Eq.2, where I_i_ is the fluorescence intensity measured at wavelength λ_i_.

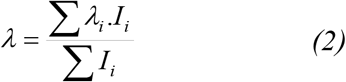

### Statistical analysis

Prior to the use of parametric tests, normality and homogeneity of variances were tested using the Shapiro-Wilk test for all quantitative data or the Levene test for raw or transformed data. First, divergences between groups were tested with one-way ANOVA, and pairwise comparisons were made with Student’s t tests. The results were expressed as the arithmetic mean for non-transformed data ± the standard error of the mean (x̅ ± SEM), except otherwise stated.

The least squares method was applied to fit functions through a regression analysis to determine the Kd values according to Eq. 1. Significance was accepted at *p* < 0.05, and Bonferroni correction was applied for sequential comparisons. All statistical analyses were performed with SPSS v. 18 for Windows.

## Acknowledgments

This work was supported by grants from INIA, MINECO, Spain to N.F.M. and E.G.F. (RTA2015-00064-C02-01 and RTA2015-00064-C02-02). The authors acknowledge financial support granted to A.V. from AGAUR (2017 SGR-229) and from the Centro de Investigación Biomédica en Red (CIBER) de Bioingeniería, Biomateriales y Nanomedicina financed by the Instituto de Salud Carlos III with assistance from the European Regional Development. We are also indebted to the CERCA Programme (Generalitat de Catalunya) and European Social Fund for supporting our research. J.V.C. received a pre-doctoral fellowship from UAB, O.C.G. received a PhD fellowship from MECD (FPU), and E.G.F. received a postdoctoral fellowship from INIA (DOC-INIA). AV has been distinguished with an ICREA ACADEMIA Award. The authors also acknowledge ICTS ‘‘NANBIOSIS”, more specifically the Protein Production Platform of CIBER in Bioengineering, Biomaterials & Nanomedicine (CIBER-BBN)/IBB, at the UAB sePBioEs scientific-technical service (http://www.nanbiosis.es/unit/u1-protein-production-platform-ppp/) and the UAB scientific-technical services LLEB, SM and SCAC (https://www.uab.cat/web/research/scientific-technical-services/all-scientific-technical-services–1345667278676.html). The authors would like to thank Milena Tileva for her helpful advice on technical issues related to the experimental adjustment of the IFN-γ activity bioassay. Special thanks to Sandra Párraga-Ferrer for the design of Fig. 3c. E. Garcia-Fruitós and N. Ferrer-Miralles designed and supervised the experiments. J.V. Carratalà, O. Cano-Garrido, J. Sánchez, C. Membrado, E. Pérez, O. Conchillo-Solé and A. Sánchez-Chardi performed the experiments. J. V. Carratalà, O. Cano-Garrido, J. Sánchez and N. Ferrer-Miralles analyzed the data. All authors edited the manuscript. N. Ferrer-Miralles wrote the paper.

